# Therapeutic *Spp1* silencing in TREM2^+^ cardiac macrophages suppresses atrial fibrillation

**DOI:** 10.1101/2024.08.10.607461

**Authors:** Noor Momin, Steffen Pabel, Arnab Rudra, Nina Kumowski, I-Hsiu Lee, Kyle Mentkowski, Masahiro Yamazoe, Laura Stengel, Charlotte G. Muse, Hana Seung, Alexandre Paccalet, Cristina Gonzalez-Correa, Emily B. Jacobs, Jana Grune, Maximilian J. Schloss, Samuel Sossalla, Gregory Wojtkiewicz, Yoshiko Iwamoto, Patrick McMullen, Richard N. Mitchell, Patrick T. Ellinor, Daniel G. Anderson, Kamila Naxerova, Matthias Nahrendorf, Maarten Hulsmans

## Abstract

Atrial fibrillation (AFib) and the risk of its lethal complications are propelled by fibrosis, which induces electrical heterogeneity and gives rise to reentry circuits. Atrial TREM2^+^ macrophages secrete osteopontin (encoded by *Spp1*), a matricellular signaling protein that engenders fibrosis and AFib. Here we show that silencing *Spp1* in TREM2^+^ cardiac macrophages with an antibody-siRNA conjugate reduces atrial fibrosis and suppresses AFib in mice, thus offering a new immunotherapy for the most common arrhythmia.

## Main text

Atrial fibrillation (AFib) is the most common irregular heart rhythm, affecting 10% of adults over 65 years old^1^. AFib leads to debilitating complications such as stroke and heart failure^2^ and, unfortunately, drugs to remedy the atrial pathologies underlying AFib remain unavailable.

Growing evidence points to the immune system^3^, specifically macrophages^4^, in the pathogenesis of AFib. In a recent study, we observed an expansion of inflammatory macrophages and atrial fibrosis in AFib patients’ atrial tissue^4^. To better dissect the relationship between macrophages, fibrosis, and AFib, we developed a mouse model that recapitulates human AFib by combining common clinical risk factors: <u>h</u>ypertension, <u>o</u>besity, and <u>m</u>itral valv<u>e</u> regurgitation (referred to as HOMER). Comparing atrial tissues from humans with AFib and HOMER mice by single cell transcriptomics documented high fidelity of the AFib animal model. We identified osteopontin, encoded by *SPP1* in humans and *Spp1* in mice, as a top upregulated gene in recruited atrial macrophages in both AFib patients and HOMER mice^4^. Osteopontin is a conserved pleiotropic matricellular protein that binds several integrins and CD44 family receptors. Macrophage-derived osteopontin stimulates fibroblasts to produce more matrix proteins and has been correlated with the progression of several chronic fibrotic diseases^5^. Fibrosis in the heart renders atrial tissue heterogeneous, which hinders homogeneous electrical conduction and acts as a structural substrate for AFib^6^. Most notably, transgenic deletion of osteopontin in monocyte-derived macrophages reduced atrial fibrosis and inducible AFib in HOMER mice^4^.

These results motivate *SPP1/Spp1* silencing in macrophages as a potential immunotherapy for AFib; however, engineering such treatment remains challenging. Although small interfering RNA (siRNA) has the potential to reversibly silence any gene and several siRNAs are under clinical development, their delivery to specific cells, especially outside the liver, remains an obstacle^7^. Currently, all five FDA-approved siRNA therapies target the liver^8^. This is largely driven by the success of conjugating siRNA to the sugar N-acetylgalactosamine (GalNAc) to elicit hepatocyte uptake. However, there is a need for delivery vehicles that target siRNAs to other cells and organs^9^. Antibodies, known for their strength and specificity in binding antigens and their established use as drug conjugates, are promising delivery vehicles for siRNA delivery^10^. Although a handful of antibody-siRNA conjugates (ARCs) targeting cancer cells have been described previously^11^, there remains an outstanding need for targeted siRNA delivery to macrophages.

Here, leveraging recent findings on the immune system’s contribution to AFib, we engineer an ARC to direct a therapeutic siRNA silencing *Spp1* to a pathogenic macrophage subset. We find that ARC therapy suppresses AFib by reducing atrial fibrosis in HOMER mice.

To achieve gene silencing, we first generated a panel of 24 predicted siRNA candidates targeting *Spp1* **(Extended Data Fig. 1)**. Based on silencing potency in murine Hepa1-6 cells and posited human cross-reactivity, we selected siRNA clone 110 (*siSpp1*) to carry forward **(Extended Data Figure 2)**. Our prior work shows human *SPP1* and mouse *Spp1* are expressed by a specific subset of atrial macrophages that upregulates triggering receptor expressed on myeloid cells 2 (TREM2) **(Fig. 1a,b)**. TREM2 is a myeloid receptor expressed on macrophages that enables phagocytosis of various ligands including phospholipids, myelin, and apoptotic cells^12^. Therefore, we selected TREM2 as an antibody target to mediate *siSpp1* delivery. We recombinantly expressed a TREM2 full-length antibody (AL002c clone 52), with previously demonstrated human and mouse TREM2 binding^13^, with an inert mouse IgG2c Fc region (**Extended Data Figure 3a**). Cellular uptake is a prerequisite for effective siRNA delivery.

**Fig. 1.**
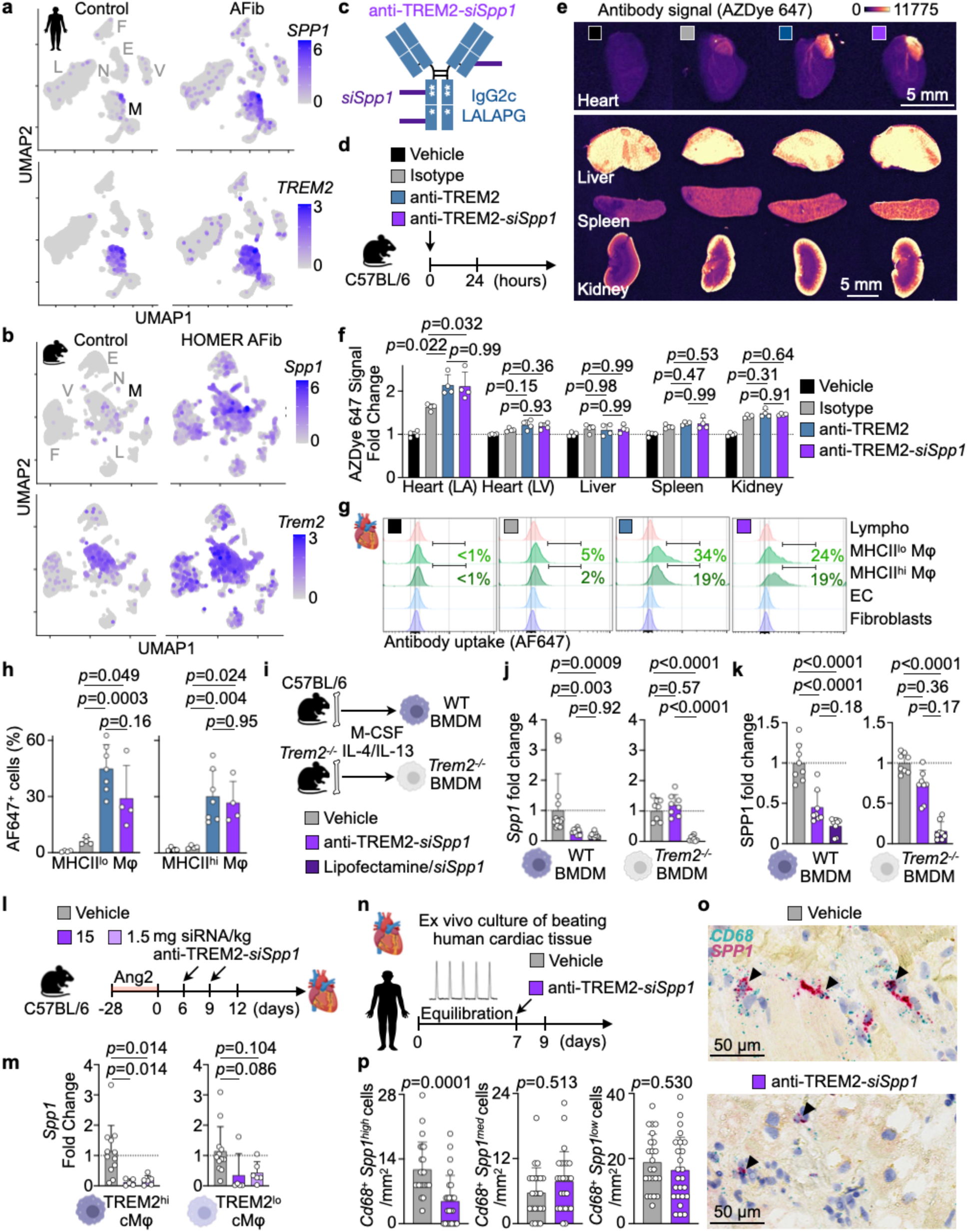
ARC silences *Spp1* in TREM2^+^ macrophages. **a**, Uniform manifold approximation and projection (UMAP) of deposited human scRNA-seq data derived from atrial appendages of five controls and seven patients with AFib highlighting co-expression of *SPP1* and *TREM2*. The major cell clusters identified were mononuclear phagocytes (M), neutrophils (N), lymphocytes (L), fibroblasts (F), endothelial cells (E), and mural cells (V). **b**, UMAP of deposited murine scRNA-seq data derived from left atria of four sham and four HOMER AFib mice highlighting co-expression of *Spp1* and *Trem2*. **c**, Schematic of a TREM2 antibody bearing an inert IgG2c Fc with LALAPG mutations (anti-TREM2) fused to siRNAs directed against *Spp1 (siSpp1)*. **d**, Experimental outline to assess the organ and cellular biodistribution of fluorescently labeled isotype control antibody, anti-TREM2, and anti-TREM2-*siSpp1* injected intravenously. **e**, Representative fluorescence images of heart, liver lobe, spleen, and kidney. Scale bars, 5 mm. **f**, Quantification of mean organ fluorescence in heart’s left atria (LA) and left ventricle (LV), liver lobe, spleen, and kidney normalized to vehicle-treated control. n = 4 per group from two independent experiments, one-way ANOVA followed by Tukey’s multiple comparisons test. **g**, Median fluorescence intensity histograms of internalized isotype control antibody, anti-TREM2, and anti-TREM2-*siSpp1* in cardiac lymphocytes (lympho), MHCII^lo^ macrophages (MHCII^lo^ Mφ), MHCII^hi^ macrophages (MHCII^hi^ Mφ), endothelial cells (EC), and fibroblasts. **h**, Percentage of total cardiac MHCII^lo^ and MHCII^hi^ macrophages with internalized fluorescence. n = 4 to 7 per group from two independent experiments, one-way ANOVA followed by Tukey’s multiple comparisons test. **i**, Experimental outline to differentiate bone-marrow-derived macrophages (BMDM) from wild-type (WT) C57BL/6 mice and *Trem2*^−/−^ mice. **j**,**k**, Quantification of *Spp1* mRNA (j) and protein (k) expression in WT (left) and *Trem2*^−/−^ (right) BMDM normalized to vehicle-treated control. n = 8 to 16 per group from three independent experiments, one-way ANOVA followed by Tukey’s multiple comparisons test. **l,** Experimental outline of C57BL/6 mice undergoing chronic hypertension for 28 days followed by treatment with vehicle or anti-TREM2-*siSpp1* on days 6 and 9. **m**, Expression of *Spp1* mRNA in TREM2^hi^ (left) and TREM2^lo^ (right) cardiac macrophage subsets isolated on day 12 normalized to vehicle-treated cells. n = 5 to 12 per group from four independent experiments, one-way ANOVA followed by Tukey’s multiple comparisons test. **n**, Experimental outline for testing anti-TREM2-*siSpp1* in beating human cardiac tissue. **o**, In situ staining of *SPP1* and *CD68* mRNA on human ventricular tissue; arrowheads indicate *SPP1^+^ CD68^+^* macrophages. **p**, Quantification of *CD68^+^* macrophages/mm^2^ with different expression levels of *SPP1* in vehicle- and anti-TREM2-*siSpp1*-treated human ventricular tissues. Vehicle: n = 23 field of views (FOV) from three heart slices; anti-TREM2-*siSpp1*: n = 25 FOV from three heart slices; nested t-test. All bar graph data are mean±s.d. with individual values for data distribution.

Therefore, we confirmed rapid internalization of anti-TREM2 by murine RAW264.7 macrophages, with only 3.99 hours required for half of the total antibody-bound TREM2 on the cell surface to internalize (**Extended Data Figure 3b-d**). Then, we successfully conjugated anti-TREM2 and *siSpp1* using click chemistry **(Fig. 1c and Extended Data Figure 4a-c)**. Size exclusion chromatography was used to remove unbound siRNA and antibody and to isolate the monomeric ARC, anti-TREM2-*siSpp1* (**Extended Data Fig. 4d**). Multi-angle light scattering analysis revealed an average siRNA-to-antibody ratio of 3 to 1 (**Extended Data Figure 4e,f**).

Next, we assessed the organ and cellular biodistribution of the ARC. Both fluorescently labeled anti-TREM2 and anti-TREM2-*siSpp1* were preferentially retained in the left atria over isotype control antibody 24 hours after intravenous injection **(Fig. 1d-f)**. No differences in drug uptake in other organs were observed **(Fig. 1e,f)**. Flow cytometry on the heart revealed internalization of anti-TREM2 and anti-TREM2-*siSpp1* by macrophages, preferentially by MHCII^lo^ macrophages **(Fig. 1g,h)**. The heart contains distinct macrophage subsets with divergent origins, functions, and prevalence in inflammation^14^. In AFib, recruited monocyte-derived macrophages accumulate in the heart^4^. We found that MHCII^lo^ macrophages, which expand in cardiac injury, expressed higher levels of the TREM2 receptor (**Extended Data Figure 5**), explaining why higher anti-TREM2 and ARC signals were detected in this macrophage subset. Negligible antibody and ARC signal could be detected in other immune cells, fibroblasts, or endothelial cells in the heart **(Fig. 1g)**, highlighting the specificity of antibody-driven uptake.

Next, we evaluated the silencing capacity of anti-TREM2-*siSpp1*. ARC exposure to wild-type C57BL/6 bone marrow-derived macrophages (BMDM) reduced *Spp1* mRNA by 70% and intracellular osteopontin protein by 50% **(Fig. 1i-k)**. The silencing achieved by antibody-mediated *siSpp1* delivery was indistinguishable from that of the gold-standard transfection reagent lipofectamine **(Fig. 1i-k)**. However, unlike lipofectamine-delivered *Spp1*, anti-TREM2-*siSpp1* did not silence *Spp1* or osteopontin in BMDM from a *Trem2*^−/−^ mouse strain, indicating that anti-TREM2-*siSpp1*’s silencing relies on binding to its target TREM2 **(Fig. 1i-k)**. Next, we confirmed target-mediated silencing in vivo using hypertensive C57BL/6 mice injected with anti-TREM2-*siSpp1* or vehicle control **(Fig. 1l,m)**. Only TREM2^hi^, but not TREM2^lo^ cardiac macrophages demonstrated a significant knockdown in *Spp1* expression by 90% at 15 mg siRNA/kg body weight dose and still 82% at a ten-fold lower dose **(Fig. 1m)**. This effect was transient; *Spp1* expression rose to 50% of baseline in TREM2^hi^ cardiac macrophages 6 days after the last dose, indicating that weekly dosing of 1.5 mg siRNA/kg body weight will be required for sustained therapeutic silencing **(Extended Data Fig. 6**). The kinetics of siRNA silencing depends on many factors including the siRNA’s potency, cellular uptake, and target mRNA or cell turnover^15^. Durable silencing of up to 6 months has been observed using GalNAc-siRNA conjugates in a clinical setting^16^. Therefore, exploring ways to prolong the silencing effect of future ARC iterations will be worthwhile.

We then evaluated the ability of anti-TREM2-*siSpp1* to silence *Spp1* in human hearts using long-term cultures of beating human ventricular myocardium from patients undergoing heart transplantation **(Fig. 1n)**. Human myocardium showed fewer *CD68^+^* macrophages with high *Spp1* expression when treated with anti-TREM2-*siSpp1* compared to vehicle control **(Fig. 1o,p)**. Importantly, anti-TREM2-*siSpp1* had no adverse effects on cardiac function and contractility of cultured human myocardium (**Extended Data Fig. 7**).

Finally and most importantly, we tested *Spp1* silencing to treat AFib. HOMER mice with established atrial disease received weekly intravenous injections of anti-TREM2-*siSpp1* or the vehicle saline **(Fig. 2a)**. After one month of treatment, these mice were subjected to an invasive electrophysiological study to examine AFib susceptibility **(Fig. 2b)**. Compared to vehicle, ARC treatment reduced AFib inducibility to 33% (from 82% in the control treated cohort) and total AFib episode burden by four-fold **(Fig. 2c)**. ARC treatment significantly reduced atrial fibrosis assayed by Masson’s trichrome staining **(Fig. 2d)**. We did not observe changes in atrial size **(Fig. 2e)**, nor liver toxicity induced by ARC treatment **(Extended Data Fig. 8)**. Collectively, these results suggest that silencing *Spp1* in TREM2^+^ macrophages is a tractable immunotherapy for AFib.

**Fig. 2.**
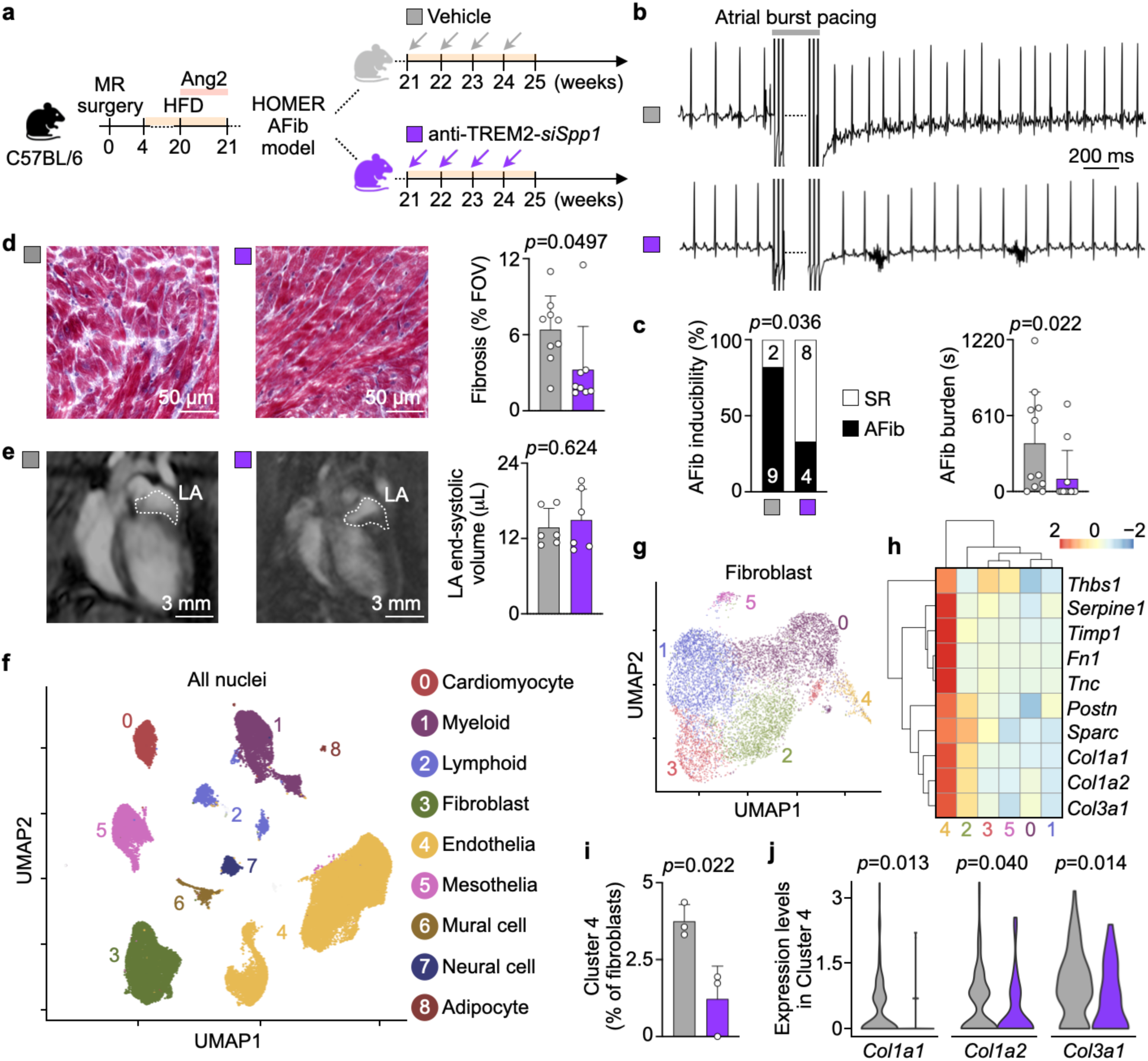
ARC therapy suppresses AFib by reducing atrial fibrosis. **a**, Experimental outline to treat HOMER mice with established atrial pathology arising from hypertension (angiotensin 2 – Ang2), obesity (high-fat diet – HFD) and mitral valve regurgitation (MR) with weekly intravenous injections of vehicle or anti-TREM2-*siSpp1* for one month. **b**, Representative ECG recordings from a cardiac electrophysiological (EP) study of vehicle- and anti-TREM2-*siSpp1*-treated HOMER mice at week 25. Scale bar, 200 ms. **c**, EP study of AFib inducibility and burden in vehicle- and anti-TREM2-*siSpp1*-treated HOMER mice. n = 11 to 12 per group from two independent experiments. (Left) Two-sided Fisher’s exact test. (Right) Two-tailed Mann– Whitney test. **d**, Representative images (left) and quantification (right) of fibrosis (Masson’s trichrome staining) in left atrial tissue from vehicle- and anti-TREM2-*siSpp1*-treated HOMER mice. n = 8 to 9 per group from two independent experiments, two-tailed Student’s t-test. Scale bars, 50 µm. **e**, Representative cardiac MRI images (left) and left atrial (LA) end-systolic volumes (right) of vehicle- and anti-TREM2-*siSpp1*-treated HOMER mice. n = 6 per group from two independent experiments, two-tailed Mann–Whitney test. Scale bars, 3 mm. **f**, UMAP delineates 9 major cardiac cell types in three vehicle- and three anti-Trem2-*siSpp1*-treated HOMER mice. Cell identities were inferred from known marker gene expression. **g**, UMAP plot of left atrial fibroblasts in three vehicle- and three anti-Trem2-*siSpp1*-treated HOMER mice. **h**, Heatmap and hierarchical clustering of the 6 fibroblast clusters according to the gene expression levels of fibrosis-related genes^46^ (represented by z-scores of the average normalized counts). **i**, Fraction of pro-fibrotic fibroblasts (cluster 4) in three vehicle- and three anti-Trem2-*siSpp1*-treated HOMER mice. Two-tailed Student’s t-test. **j**, Expression levels of collagen genes in pro-fibrotic fibroblasts (cluster 4) of three vehicle- and three anti-Trem2-*siSpp1*-treated HOMER mice. Kolmogorov-Smirnov test. All bar graph data are mean±s.d. with individual values for data distribution.

To interrogate the pathways leading to the observed treatment benefit, we collected left atria from HOMER mice after anti-TREM2-*siSpp1* or vehicle treatment and performed single nuclei RNA sequencing (snRNA-seq) using the 10X Genomics Chromium platform. A total of 48,647 single nuclei were partitioned into clusters **(Fig. 2f)**. We identified 8 noncardiomyocyte cell types: myeloid, lymphoid, fibroblasts, endothelial, mesothelia, mural cells, neural cells, and adipocytes **(Fig. 2f and Extended Data Fig. 9a,b)**. Given the reduction of atrial fibrosis after ARC treatment, we closely examined the fibroblasts **(Fig. 2g)**. All fibroblast subclusters express matrix-related genes to varying degrees, but we observed that *Col1a1, Col1a2, Col3a1,* and others were most expressed in cluster 4 **(Fig. 2h)**. Cluster 4 fibroblasts also exclusively expressed *Cd44* **(Extended Data Fig. 9c**), which we previously identified as a potential SPP1 receptor in atrial fibroblasts from HOMER mice and AFib patients^4^. Cluster 4 fibroblasts were reduced in the ARC-treated group compared to the vehicle arm **(Fig. 2i)**. Furthermore, the expression of several collagen genes in cluster 4 cells was significantly lowered by ARC treatment **(Fig. 2j)**. Although cluster 4 comprises <5% of total fibroblasts, their small proportion is consistent with the notion that a niche of cells can regulate global tissue changes during disease^17,18^. Reduction of both matrix-producing cells **(Fig. 2i)** and genes **(Fig. 2j)** aligns with less fibrosis **(Fig. 2d)** and suppression of AFib **(Fig. 2b,c)** achieved by ARC therapy.

To our knowledge, our study provides the first demonstration of antibody-leveraged siRNA delivery to a specific subset of pathogenic macrophages. While we and others have previously used lipid nanoparticles to deliver siRNA to cardiovascular macrophages^19,20^, nanoparticles were not directed to a disease-promoting macrophage subset, and uptake by other cell types and tissues remained an issue. ARCs, as evidenced by anti-TREM2-*siSpp1*, are suited to overcome biodistribution challenges and restrict siRNA uptake and gene silencing to specific cell subsets within a tissue. Efficient silencing of *Spp1* in TREM2^+^ macrophages may provide a means to treat other fibrosis-driven pathologies such as fatty liver disease^21^, idiopathic pulmonary fibrosis^22^, systemic sclerosis^23^, and chronic kidney disease^24^.

While no immunotherapies have been approved for treating AFib, modest effects have been observed in small trials using colchicine^25,26^, an anti-inflammatory small molecule, and canakinumab^27^, an IL-1β neutralizing antibody. Unfortunately, these agents also disrupt essential immune functions in healthy tissues, leading to an increased risk of infections which limits their chronic clinical use^28–30^. A daunting challenge in the development of safe and effective immunotherapies is ensuring their effects are isolated and focused on the target cells and diseased tissue^31,32^. By conferring cell specificity to gene silencing, ARCs emerge as a promising immunotherapy modality. Specifically, our study using anti-TREM2-*siSpp1* alongside other recent efforts^33^ showcases the feasibility of targeted immunotherapy to treat AFib.

Although silencing *Spp1* in TREM2^+^ cardiac macrophages suppressed AFib in a majority of HOMER mice, a subset of treated mice still experienced short episodes of AFib. Likely, other pathways driving AFib remain active. Engineering new ARCs targeting other genes and cells, including cardiomyocytes and fibroblasts, may be an option to treat the multifaceted etiology of AFib in heterogenous patient subsets.

## Acknowledgments

The authors thank the HSCI-CRM Flow Cytometry Core for assistance with cell sorting, and scientists at Axolabs for assistance in generating, validating, and producing siRNA used in these studies. We acknowledge BioRender (IH23GLWC6I) for cartoon components.

## Author Contributions

N.M. conceived the study; designed, performed, and analyzed all experiments; interpreted data and made figures. S.P. designed, performed, and analyzed experiments; interpreted data and made figures. A.R., M.Y., M.J.S., S.S., and R.N.M. provided critical reagents and resources. N.K., K.M., L.S., C.G.M., H.S., A.P., C.G.C., E.B.J., J.G., G.W., Y.I., and P.M. designed, performed, and analyzed experiments. I.L. and M.H. analyzed snRNA-seq data and made figures. P.T.E., D.G.A, K.N., M.N., and M.H. discussed results and strategy. M.N. and M.H. conceived and directed the study. N.M., S.P., M.N., and M.H. wrote the manuscript with input from all the authors.

## Data and materials availability

The authors declare that all data supporting the findings are available within the paper. Mouse snRNA-seq data are available at the NCBI’s Gene Expression Omnibus database under accession no. GSE272503.

## Materials and Methods

### Humans

All procedures were performed according to the Declaration of Helsinki and were approved by the local ethics committee of the University of Regensburg (20-1776-101), University of Göttingen (31/9/00) and by the Mass General Brigham Institutional Review Board (2023P002856). Informed consent was obtained from all patients. Left-ventricular myocardium was acquired from explanted hearts of patients with end-stage heart failure undergoing heart transplantation. Myocardial tissue was immediately placed into cooled cardioplegic solution (4°C) after excision for transportation.

### Mice

C57BL/6 (stock 000664), and C57BL/6J-*Trem2^em2Adiuj^*/J (*Trem2*^−/−^; stock 027197) mice were purchased from The Jackson Laboratory. All experiments were performed with 8-to-12-week-old male animals and using age-matched control groups. Where appropriate, mice were randomly assigned to groups. Housing facilities followed a 12-hour dark/light cycle, room temperature was 18–22°C, and maintained humidity was between 40% and 60%. Mice were fed with respective diets ad libitum. All animal experiments were approved by the Institutional Animal Care and Use Committee at the Massachusetts General Hospital (2014N000078) in accordance with federal, state and local guidelines.

### HOMER model of inducible AFib

The HOMER mouse model, described previously to induce atrial pathology by combining hypertension, obesity, and mitral valve regurgitation^4^, was used to assess the therapeutic efficacy of *Spp1* silencing on AFib inducibility. In brief, mice underwent surgery to generate mitral valve regurgitation. Four weeks after surgery, high-fat diet (Research Diets, D12492) was initiated and continued for at least 17 weeks. Finally, immediately prior to treatment initiation, mice were exposed to continuous infusion of Ang2 (1 μg/kg/min) for one week via a subcutaneously implanted osmotic minipump (Alzet). These HOMER mice were treated with a total of four weekly intravenous injections of either vehicle (100 µL of PBS) or anti-TREM2-*siSpp1* (1.5 mg siRNA/kg mouse weight in 100 µL of PBS). During the course of treatment, HOMER mice remained on a high-fat diet.

### Hypertension model to assess *Spp1* silencing

The expression of *Spp1* is low in a healthy naive animal (**Fig. 1b**). Therefore, we induced hypertension, described previously to upregulate *Spp1* expression in the heart^34^ to assess the in vivo silencing of anti-TREM2-*siSpp1* in a high-throughput manner. Ang2 (1 µg/kg/min) was continuously administered through osmotic minipumps (Alzet) for 28 days after which they were removed. After a 6-day recovery period, mice were randomized into treatment groups and intravenously injected with a total of 15 or 1.5 mg siRNA/kg body weight of anti-TREM2-*siSpp1* or vehicle (PBS) divided into two consecutive doses, three days apart. Hearts were excised three days or six days after the last injection.

### Organ and cellular biodistribution

For organ biodistribution assessments, 8-week-old male C57BL/6 mice were injected intravenously with 100 µL of PBS vehicle or 1 nmol of AZDye 647-labeled isotype antibody, anti-TREM2, or anti-TREM2-*siSpp1* in an equivalent volume. Exactly 24 hours later, 20 mL of ice-cold PBS was used to perfuse the mouse via the left ventricle. The heart, spleen, kidney, and liver lobe were excised and collectively imaged on an Azure Sapphire FL Biomolecular Imager.

For cellular biodistribution assessments, 8-week-old male C57BL/6 mice were injected intravenously with 100 µL of PBS vehicle or 1 nmol of AF647-labeled isotype antibody, anti-TREM2, or anti-TREM2-*siSpp1* in an equivalent volume. Exactly 24 hours later, 20 mL of ice-cold PBS was used to perfuse the mouse via the left ventricle. The heart was processed for flow cytometry as described below; however, TREM2-APC antibody was excluded from the staining panel.

### Electrophysiological study

Electrophysiological studies were performed as previously described by us^4^. Mice were anesthetized with isoflurane and temperature controlled using a homeothermic monitoring system (Harvard Apparatus). A Millar EPR-800 octapolar catheter was inserted into the right jugular vein and positioned in the right atrium and ventricle. AFib inducibility was tested via atrial burst pacing for three seconds with decremental cycle lengths starting at 50 ms and progressing to 10 ms in intervals of 5 ms. The protocol was repeated with a burst pacing duration of 6 ms. AFib was defined as irregular ventricular response with rapid atrial rhythm. Mice were considered inducible if they had at least one episode of AFib longer than one second.

### Ex vivo culture of beating human cardiac tissue

Human ventricular tissue was cut in 300 µm slices in 4°C slicing buffer using a vibratome. Myocardial slices were attached to plastic triangles and mounted in biomimetic culture chambers with an integrated magnetic force sensor and pacing electrodes^35^. A mechanical preload was introduced, and electric culture pacing was started. Beating myocardium was cultured at 37°C and 5% CO_2_ on a rocking platform using M199 medium, supplemented with 1% ITS, 1% Pen/Strep, 50 µM β-ME and 20 nM hydrocortisone. Medium was changed every 2-3 days. After 7-10 days of equilibration, beating myocardial slices (60 bpm) were treated with anti-TREM2-*siSpp1* or vehicle control for 48 hours. Contractility, force-frequency relationship and post-rest behavior were assessed before formalin fixing and paraffin embedding the tissue for RNA in situ hybridization.

### Cell lines

Expi293 cells were obtained from Thermo Fisher Scientific and maintained in Expi293 Expression medium. RAW264.7 cells were obtained from the American Type Culture Collection and maintained in Dulbecco’s modified Eagle medium (DMEM) enriched with 10% (v/v) heat-inactivated FBS. All cells tested negative for mycoplasma and were maintained at 37°C and 5% CO_2_.

### Bone marrow-derived macrophage differentiation and cell assays

Femurs from C57BL/6 or *Trem2*^−/−^ mice were flushed and filtered to isolate bone marrow cells. These cells were cultured in DMEM enriched with 10% (v/v) heat-inactivated FBS supplemented with 30 ng/mL of M-CSF. After 5 days of culture, bone-marrow derived macrophages (BMDM) were polarized by adding 10 ng/mL of IL-4 and IL-13 to the medium for 24 hours. BMDM were treated with either vehicle OptiMEM (Thermo Fisher Scientific), anti-TREM2-*siSpp1,* or *siSpp1* complexed with lipofectamine RNAiMAX transfection reagent (Thermo Fisher Scientific) for 48 hours.

### siRNA identification, validation and development

With assistance from Axolabs, we employed a bioinformatic approach to identify a screening set of 24 siRNA candidates directed against mouse *Spp1* mRNA (secreted phosphoprotein 1, NCBI Gene ID 20750). This approach assumes a canonical 19-mer dsRNA structure whereby positions 2-18 (5’-3’) of the sense and antisense strand were used for specificity calculations. Positions 2-7 (5’-3’) were considered the seed region and used to exclude candidates with identical seed regions to known miRNAs. Positions 1-19 (5’-3’) of the antisense strand were used for human cross-reactivity analysis; perfect matches or single nucleotide mismatches were considered cross-reactive hits.

Candidate siRNAs were synthesized and purified with RNA, DNA, 2’-O-methyl residues, and phosphorothioate modifications at defined positions. Each single strand was HPLC purified to >85% purity and its mass confirmed by LC/MS. Candidate siRNA’s specific silencing was validated in murine Hepa1-6 cells using lipofectamine as a transfection reagent and after 24 hours of exposure. *Spp1* and *Gapdh* mRNA were measured by branched DNA assay. siRNA against *F-luc*, *Ahsa1*, and *FVII* were used as negative controls.

For the siRNA candidate selected for antibody conjugation, the siRNA sense and antisense strands were generated using the standard pattern of alternating 2’-O-methyl and 2’-fluoro nucleotides and phosphorothioate bonds at the extremities of the strands^36^. For the anti-sense strand, a 5’-(E)-vinylphosphonate (5’-E-VP) 2’-O-methyl uridine was included as it was reported that 5’-E-VP has an increased fit inside an Ago2-5’-phosphate binding pocket relative to the native strand and its metabolic stability increases tissue accumulation^37^. For the sense strand, an aminohexyl linker was appended to the 3’ end to facilitate the addition of a pegylated trans-cyclooctene (TCO) group for downstream chemical conjugation using TCO-Peg4-NHS from Broadpharm^38^.

### Antibody production

Plasmids encoding the variable heavy and light chain of a human/mouse cross-reactive antibody directed against TREM2 clone 52 were derived from a patent application (WO 2016/023019 A2). The variable heavy chain was fused to mouse IgG2c fragment crystallizable region (Fc) bearing LALAPG mutations rendering the Fc region inert. The plasmids were transfected into Expi293 cells following manufacturer’s recommendations (Thermo Fisher Scientific). The antibodies produced by these cells were isolated by protein A Sepharose affinity chromatography. Antibody was then further purified by size exclusion chromatography using a HiLoad 16/600 Superdex 200 pg column on an ÄKTA FPLC system (GE Healthcare) that had been pretreated for 4 hours with 1 M NaOH to remove endotoxin and subsequently equilibrated in sterile PBS (Corning). After purification, all proteins were buffer exchanged into sterile PBS, 0.2 µm sterile-filtered (Pall Corporation), and confirmed to contain minimal endotoxin (<0.1 EU/mL) using a chromogenic LAL assay (Lonza).

### Antibody-siRNA conjugation and validation

To conjugate the siRNA to the antibody, we employed a click chemistry-based strategy^38^. Lysine residues on the antibody were coupled with pegylated methyl tetrazine (MeTz) using the corresponding NHS ester and covalently linked to a TCO handle on the 3’ end of the sense strand of the hybridized siRNA at a ratio of 1:10. ARCs were subsequently purified by size exclusion chromatography, removing uncoupled siRNA and antibody. The RNA content was quantified by UV or ribogreen assay and the protein content by BCA assay (both Thermo Fisher Scientific) according to the manufacturer’s protocol.

### SEC-MALS method for MW and drug-to-antibody ratio determination

Size exclusion chromatography with multi-angle light scattering (SEC-MALS) data collection was performed on an analytical system consisting of an Agilent Technologies 1260 Infinity II HPLC system (sampler, pump, and UV–vis detector) equipped with a Wyatt Technologies Heleos II multi-angle light scattering detector with in-line QELS detector for DLS and Wyatt Technologies Optilab T-rEX refractive index detector, using an analytical SEPAX SRT SEC-300 SEC (300 Å, 2.7 μm, 7.8 × 300 mm) column. PBS 1× (137 mM NaCl, 2.7 mM KCl, 10 mM phosphate, pH 7.0) as the mobile phase eluting at a rate of 0.5 mL min^−1^ and run time of 45 minutes was used for each experiment (injected sample concentrations and volumes: 0.7 mg mL^−1^ in 100 μL of PBS for anti-TREM2 and 0.353 mg mL^−1^ in 100 μL of PBS for anti-TREM2-*siSpp1*). Molecular weights were determined using a refractive index increment (dn/dc) of 0.1700 mL/g.

### Fluorophore labeling

Antibodies or ARC were labeled with Alexa Fluor 647 (AF647) N-hydroxysuccinimide ester, Alexa Fluor 488 (AF488) N-hydroxysuccinimide ester (both Thermo Fisher Scientific), or AZDye 647 Hydrazide (Vector Labs). Excess dye was removed using a PD10 desalting column (GE Healthcare), and the degree of labeling for each protein was calculated. Anti-TREM2 labeled with AF488 was used to confirm TREM2 binding and measure net internalization rate. Antibodies and ARC injected for organ biodistribution assays possessed equimolar AZDye 647, 1 nmol. Antibodies and ARC injected for cellular biodistribution assays possessed equimolar AF647, 1 nmol.

### Anti-TREM2 binding confirmation

Expi293 cells or RAW264.7 cells in suspension were incubated with 100 nM of anti-TREM2-AF647 in FACS buffer (PBS containing 0.5% w/v of bovine serum albumin and 2 mM of EDTA) on ice for 30 minutes. After treatment, cells were washed and antibody binding was measured by flow cytometry on an LSRII (BD Biosciences) and analyzed with FlowJo software.

### Antibody net internalization assay

RAW264.7 cells were treated with 100 nM anti-TREM2-AF488 for time points between 0–5 hours. Based on the dissociation and association rates, this concentration range ensured a rapid equilibration rate, with the resulting equilibrium favoring saturated surface receptors. After treatment, cells were washed once with PBS and then detached from the plate using 0.25% Trypsin/EDTA. Cells were pelleted at 1000×*g* for 5 minutes and then resuspended in FACS buffer.

To determine what fraction of the total anti-TREM2-AF488 signal derived from surface-bound antibody rather than internalized antibody, we used an anti-AF488 polyclonal antibody (Thermo Fisher Scientific) which exhibits >90% quenching efficiency. Half of the pelleted sample was diverted for staining with 20 µg/mL of anti-AF488, followed by incubation of both samples on ice for 10 minutes. The mean fluorescence intensity (MFI) of AF488 signal for all samples (quenched versus unquenched, different timepoints, and replicates) was measured via flow cytometry on an LSRII and analyzed with FlowJo software.

To calculate the internalized fraction, we assumed that the difference in MFI between the unquenched and quenched fixed cells arises from cell surface-bound antibody. The internalized fraction was then determined by subtracting the cell surface-bound antibody AF488 MFI from total unquenched cell AF488 MFI. Besides the initial time point, the surface TREM2 receptors were considered fully saturated. Assuming internalization is a first-order rate process, we fitted a linear relationship between the internalized fraction and the surface integral to determine the net internalization rate.

### Flow cytometry cell preparation from heart tissue

Mice were perfused through the left ventricle with 20 mL of ice-cold PBS. Hearts were then excised and left atrial tissues were microdissected using a dissection microscope. After harvest, tissues were minced into small (1 mm^3^) pieces using scissors and enzymatically digested with 450 U/mL of collagenase I, 125 U/mL of collagenase XI, 60 U/mL of DNase I, and 60 U/mL of hyaluronidase (all Sigma-Aldrich) for 60 minutes at 37°C under agitation. Tissue pieces were mechanically dissociated through a 40 μm nylon mesh filter, washed with ice-cold FACS buffer and then centrifuged (340×g for 7 minutes) to obtain single-cell suspensions ready for staining.

### Flow cytometry staining, acquisition, and analysis

To detect cellular subsets in the heart, cell suspensions were stained at 4°C in FACS buffer. Heart single cell suspensions were first stained with Zombie Violet viability dye (BioLegend) and blocked with CD16/CD32 antibody (clone 93, 1:100, eBioscience). Cells were then stained with CD45-BV711 (clone 30-F11, 1;600, BioLegend), CD3-BV785 (clone 17A2, 1:100, BioLegend), CD19-BV785 (clone 6D5, 1:100, BioLegend), Ly6G-FITC (clone 1A8, 1:600, BioLegend), CD31-PerCP/Cy5.5 (clone 390, 1:400, BioLegend), MEFSK4-PE (clone mEF-SK4, 1:100, Miltenyi Biotec), CD64-PE/Cy7 (clone X54-5/7.1, 1:250, BioLegend), MHCII-AF700 (clone M5/114.15.2, 1:600, BioLegend), CD11b-APC/Cy7 (clone M1/70, 1:600, BioLegend), and TREM2-APC (clone 237920, 1:20, R&D Biosystems). Data were acquired on an Attune NxT (Thermo Fisher Scientific) and analyzed with FlowJo software.

### Flow cytometry gating

Heart cell subsets were defined based on marker expression as follows: endothelial cells as CD45^−^CD31^+^, fibroblasts as CD45^−^CD31^−^MEFSK4^+^, neutrophils as CD45^+^CD11b^+^Ly6G^+^, lymphocytes as CD45^+^CD11b^−^CD3^+^ and CD45^+^CD11b^−^CD19^+^, and cardiac macrophages as CD45^+^CD11b^+^CD64^+^.Cardiac macrophages were further parsed by MHCII expression. The gating strategy employed is shown in **Extended Data Fig. 5b**.

### FACS isolation of cardiac macrophages

Cardiac cell suspensions were stained as described above. Macrophages were sorted according to their MHCII expression on a FACSAria II cell sorter (BD Biosciences) directly into 350 µL of RLT Plus lysis buffer (Qiagen).

### Real-time qPCR

Total RNA was extracted from BMDM using the AllPrep RNA/Protein kit or from FACS-purified macrophages using the RNeasy Plus Micro kit (both Qiagen) according to the manufacturer’s protocol. RNA was reverse transcribed using the High-Capacity RNA-to-cDNA kit (Thermo Fisher Scientific) according to the manufacturer’s instructions. TaqMan Fast Universal PCR Master Mix (Thermo Fisher Scientific) and TaqMan gene expression assays for *Spp1* (Mm00436767_m1, FAM-MGB) and *Gapdh* (Mm99999915_g1, VIC-MGB, both Thermo Fisher Scientific) were used to quantify *Spp1* mRNA. The relative changes were normalized to *Gapdh* mRNA using the 2^−ΔΔCt^ method.

### Osteopontin ELISA

Total protein extracted from BMDM using the AllPrep RNA/Protein kit (Qiagen) was analyzed with the Osteopontin ELISA kit (Abcam) according to the manufacturer’s protocol.

### Histology

To detect left atrial fibrosis, hearts were embedded in OCT compound and fresh-frozen sections were prepared for Masson’s Trichrome staining. Sections were fixed in 10% formalin for 30 minutes at room temperature and incubated in Bouin’s fixative solution (Electron Microscopy Sciences) overnight at room temperature. The nuclei were stained with a mixture of Weigert’s iron hematoxylin solution A and B (Electron Microscopy Sciences), followed by staining with Biebrich scarlet-acid fuchsin (StatLab). After the sections were incubated in phosphomolybdic acid/phosphotungstic acid solution (Sigma-Aldrich), collagen was stained with aniline blue stain (StatLab). All slides were scanned by a digital slide scanner NanoZoomer 2.0RS (Hamamatsu). To analyze fibrosis from stained tissue sections, ten fields of views (FOV) from the left atrium per animal were randomly selected by a person blinded to cohort assignment and exported at 40× magnification using NanoZoomer NDP.view2 software (Hamamatsu). For each FOV, the percentage of positive signal was measured using ImageJ v2.9.0 (NIH) software.

To assess hepatic histological features, livers were submerged in 15 mL of 10% neutral buffered formalin solution (Sigma) overnight at room temperature. Livers were then placed in 70% ethanol prior to paraffin embedding and processing. Next, paraffin-embedded livers were sectioned at 5 µm thickness and stained with hematoxylin (Polysciences) and eosin (Sigma-Aldrich). Stained liver sections were assessed and scored by a blinded, trained pathologist according to the degree of steatosis, inflammation, and necrosis on a 0-4 severity scale (in which 0 corresponds to no hepatocyte lipid accumulation, no inflammation, and no necrosis).

### Human RNA in situ hybridization

In situ hybridization was performed on a Leica Biosystems BOND RX system using human FFPE samples and the RNAscope 2.5 LS Duplex assay with RNAscope 2.5 LS Green Accessory pack (both Advanced Cell Diagnostics) following the manufacturer’s protocol with minor modifications. These modifications include an antigen retrieval step for 15 minutes at 88°C and a protease incubation step for 15 minutes at 40°C. Each FFPE sample was first stained with the RNAscope 2.5 LS Duplex positive (Hs-PPIB-C1, Hs-POLR2A-C2) and negative (DapB-C1, DapB-C2) control probes to assess tissue quality. If the sample passed quality control metrics (i.e., sufficient positive control probe signal in the absence of negative control probe signal), the adjacent tissue section was stained with the RNAscope 2.5 LS Hs-CD68-C1 (Cat# 560598) and Hs-SPP1-C2 (Cat# 420108-C2) probes to detect *CD68* and *SPP1* signal, respectively. For quantification, *SPP1^+^*dots were manually counted using ImageJ software and *SPP1^+^CD68^+^*macrophages were binned based on the number of *SPP1^+^* dots per cell. *SPP1^low^* macrophages were characterized by 0-3 dots/cell, *SPP1^med^* macrophages by 3-6 dots/cell, and *SPP1^high^*macrophages by >6 dots/cell.

### Cardiac MRI

MRI was performed with a 4.7-Tesla horizontal bore Pharmascan system (Bruker) equipped with a custom-built mouse cardiac coil (Rapid Biomedical) to obtain cine images of the left atrial and ventricular short axes and a cine loop of the horizontal long-axis. ECG and respiratory gating (SA Instruments), a gradient echo FLASHsequence and a dedicated mouse cardiac volume coil in bird cage design were used. Imaging parameters were as follows: echo time (TE), 2.7 to 5 ms; 16 frames per RR interval (TR 7.0-15 ms depending on heart rate); in-plane resolution, 150×150 µm; slice thickness, 1 mm; NEX 4; flip angle, 30 degrees. Left atrial size was measured using Horos software.

### snRNA-seq

For snRNA-seq experiments, left atrial tissues from vehicle- and anti-TREM2-*siSpp1*-treated HOMER mice were microdissected using a dissection microscope and subsequently snap-frozen in liquid nitrogen. Nuclei were then isolated using the 10X Chromium Nuclei Isolation kit according to the manufacturer’s protocol. In detail, frozen samples were homogenized using a pestle and lysis buffer and passed through a column. Next, debris was removed via centrifugation in debris removal buffer. The nuclei were then washed, resuspended and visualized under a microscope to determine nuclei concentration and quality. The isolated nuclei were immediately loaded on a 10X Genomics Chromium Single Cell G chip as outlined by the 10X Genomics Chromium Single Cell 3’ Reagent kit v3.1 user guide. As nuclei have a lower RNA content than whole cells, the cDNA amplification cycles were increased by 1-2 cycles, as recommended by 10X Genomics. Libraries were then prepared according to the manufacturer’s protocol with modifications for PCR cycles based on the cDNA concentration. Constructed libraries were validated and quantified with the Agilent High Sensitivity DNA chip and by real-time qPCR and subsequently sequenced on an Illumina NovaSeq 6000 system as recommended by 10X Genomics.

### scRNA-seq and snRNA-seq analysis

We analyzed deposited scRNA-seq data (GSE224959) as previously described by us^4^. For snRNA-seq data, the sequencing reads were aligned to the mouse (mm10) reference genome and processed with 10X Genomics Cell Ranger v7.1.0, which generated a gene-cell barcode count matrix for each sample. These count matrices were modified by removal of ambient RNA contamination using CellBender v0.3.0^39^. Quality-control metrics, including the number of unique molecular identifiers (nUMI) detected, the number of unique genes (nGene) detected, and the percentage of reads mapped to the mitochondrial genes (percent.mt), were used to filter out low-quality nuclei. Specifically, for each sample, nuclei were filtered out if their nUMI or nGene were lower than their respective sample median by more than three times the median absolute deviation (MAD), or if their percent.mt were higher than the sample median by more than three MADs. Further hard cutoffs of nUMI > 150, nGene > 150, and percent.mt < 5 are imposed across the samples. Doublets were flagged with scDblFinder^40^. Seurat v4.3.0^41–43^ was used to carry out further analyses, including count normalization with sctransform v2^44,45^ followed by an integration workflow to identify shared cell states present across all samples under consideration, as well as nuclei clustering and cluster marker identification. Clusters with more than 40% of nuclei flagged as doublets were removed, and the same Seurat analyses were repeated with the remaining nuclei.

### Statistics and reproducibility

Statistical analyses were performed using GraphPad Prism 9. Normality of data was assessed using the Kolmogorov Smirnov test. The specific statistical methods and number of replicates are indicated in the figure legends. *p*-values of 0.05 or less were considered significant. Animal group sizes were mindfully and empirically chosen. Animals were randomly assigned to treatment groups. Data were analyzed in a blinded fashion where appropriate.

---

**Extended Data Fig. 1.**
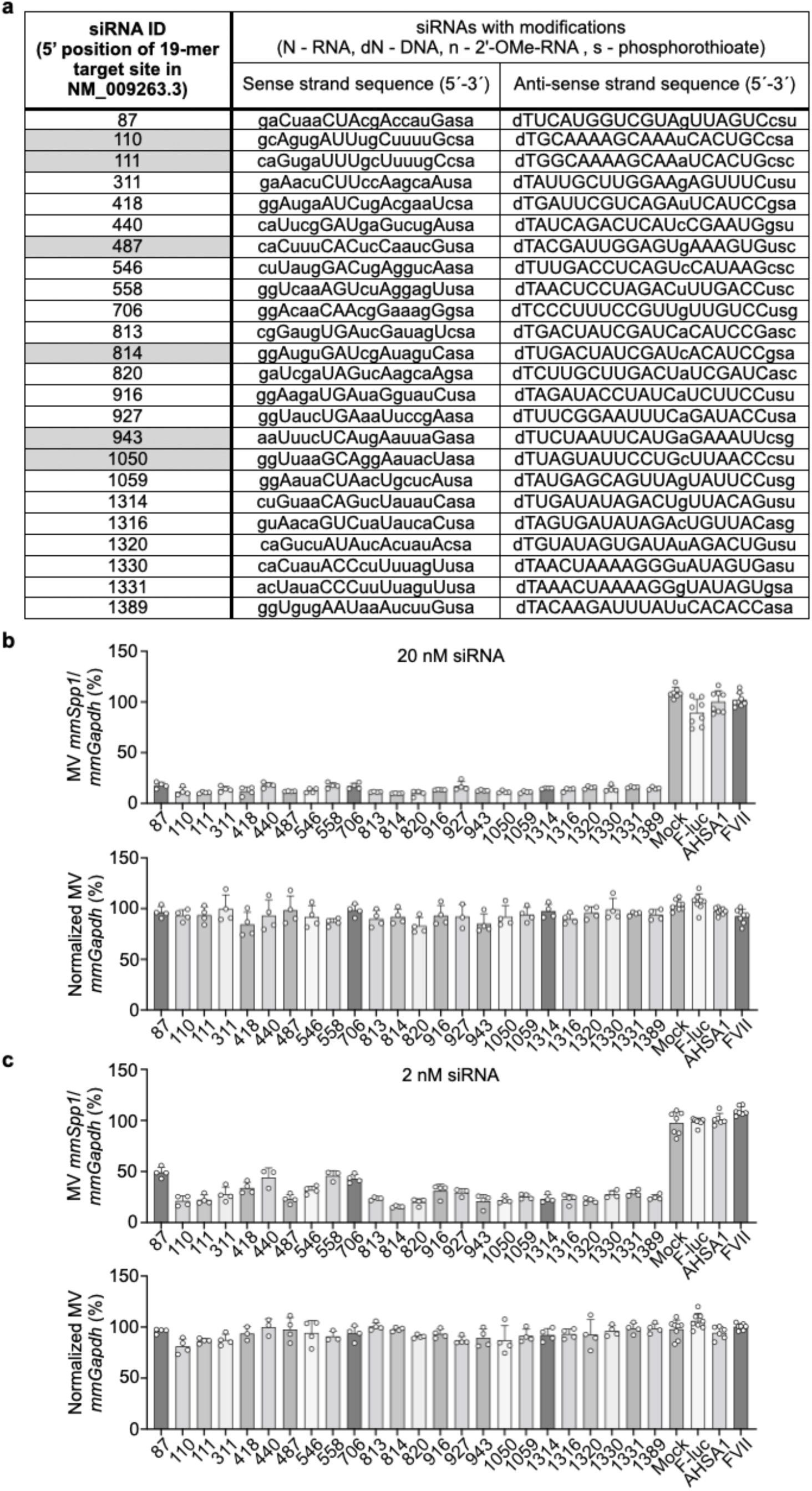
Dual dose response of predicted siRNA candidates. **a**, Predicted siRNA sequences directed against mouse *Spp1* (*mmSpp1*). Gray boxes denote siRNA candidates with predicted cross-reactivity to human *SPP1*. **b**,**c**, The effect of each candidate siRNA in Hepa1-6 cells at 20 nM (b) and 2 nM (c) concentrations on target *mmSpp1* expression normalized to *mmGapdh* expression (top) and absolute *mmGapdh* expression normalized to mock-transfected cells (bottom). n = 4 to 8 biological replicates per candidate. All bar graph data are mean±s.d. with individual values for data distribution.

**Extended Data Fig. 2.**
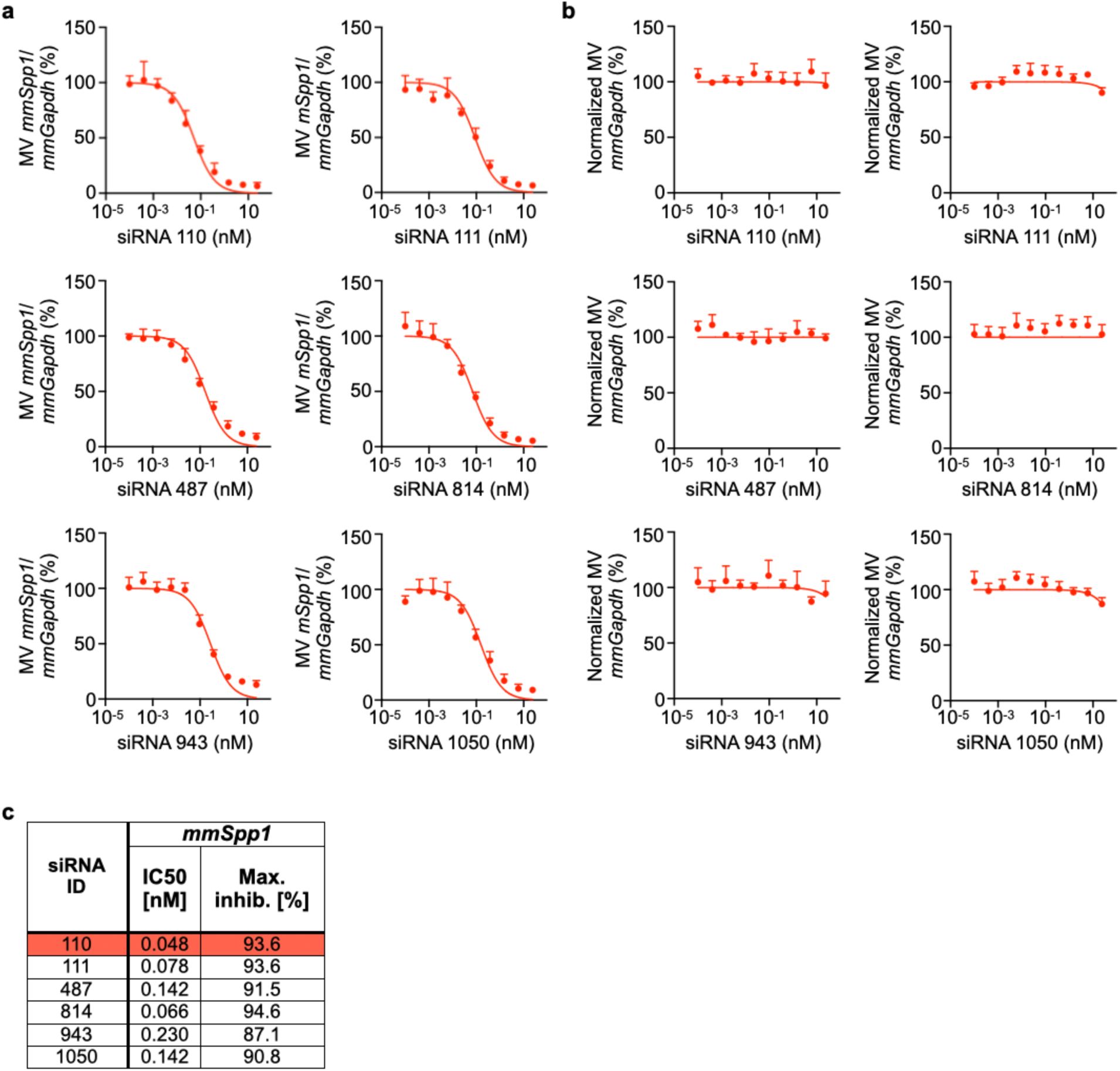
Dose response of top siRNA candidates. **a,b**, The effect of siRNA candidates 110, 111, 487, 814, 943, and 1050 in Hepa1-6 cells at different doses on target *mmSpp1* expression normalized to *mmGapdh* expression (a) and absolute *mmGapdh* expression normalized to mock-transfected cells (b). Data are mean±s.d., n = 4 biological replicates per dose per candidate. **c**, The silencing potency (IC50) and maximum inhibition (max. inhib.) of top siRNA candidates generated from a sigmoidal dose-response curve fit.

**Extended Data Fig. 3.**
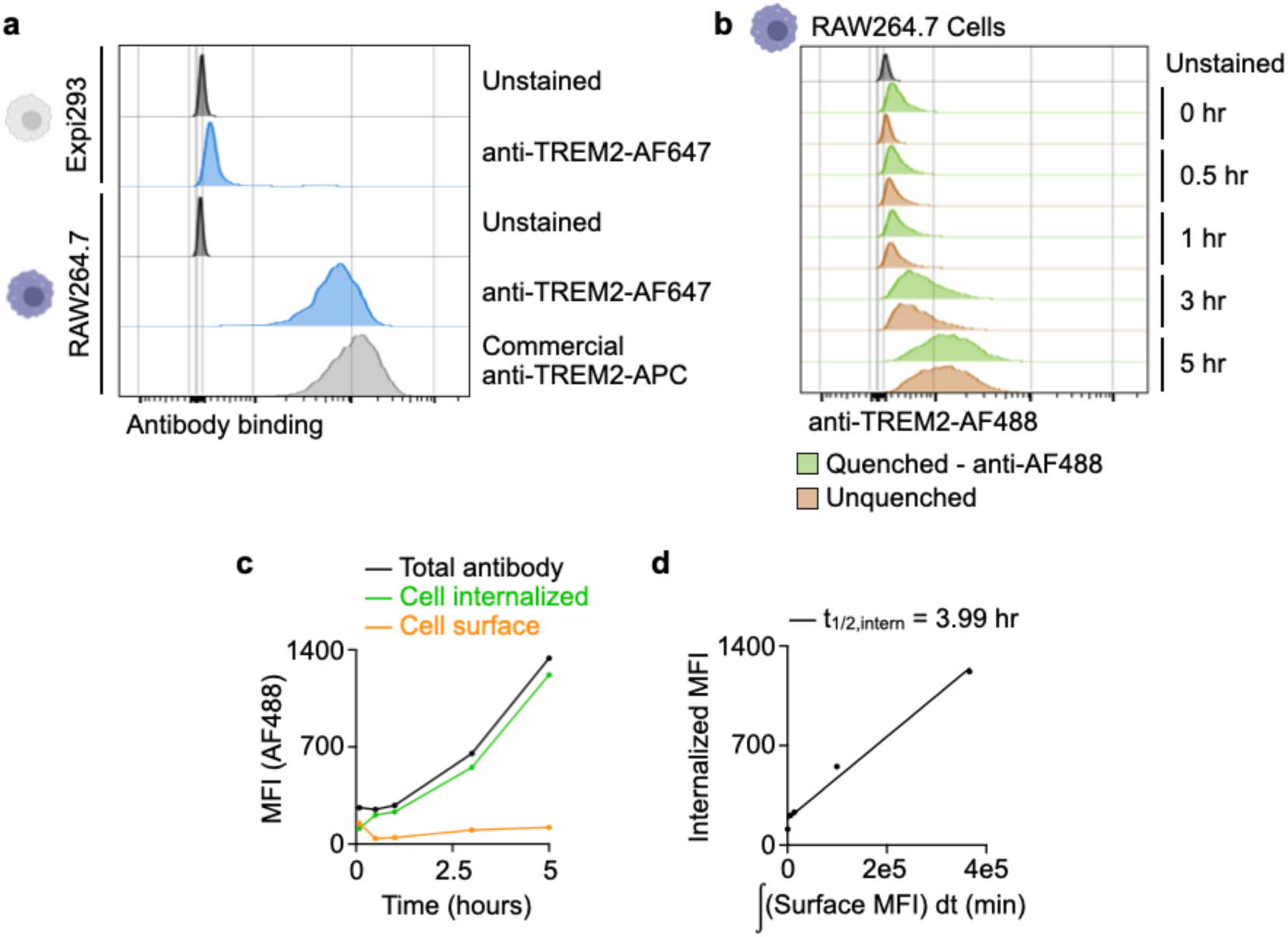
Binding and internalization of anti-TREM2. **a**, Antibody binding of AF647-conjugated anti-TREM2 in RAW264.7 macrophages by flow cytometry. Expi293 cells lacking TREM2 expression served as negative control cells, while commercially available TREM2-APC antibody served as a positive control antibody. **b**, RAW264.7 macrophages exposed to AF488-conjugated anti-TREM2 for 0, 0.5, 1, 3, and 5 hours with or without exposure to the anti-AF488 fluorescence quenching antibody. **c**, MFI quantification attributed to cell surface-binding antibody, cell internalized antibody, and total antibody signal. **d**, Linear fit of the internalization rate using measurements of the internalized fraction of antibody versus the surface integral over time.

**Extended Data Fig. 4.**
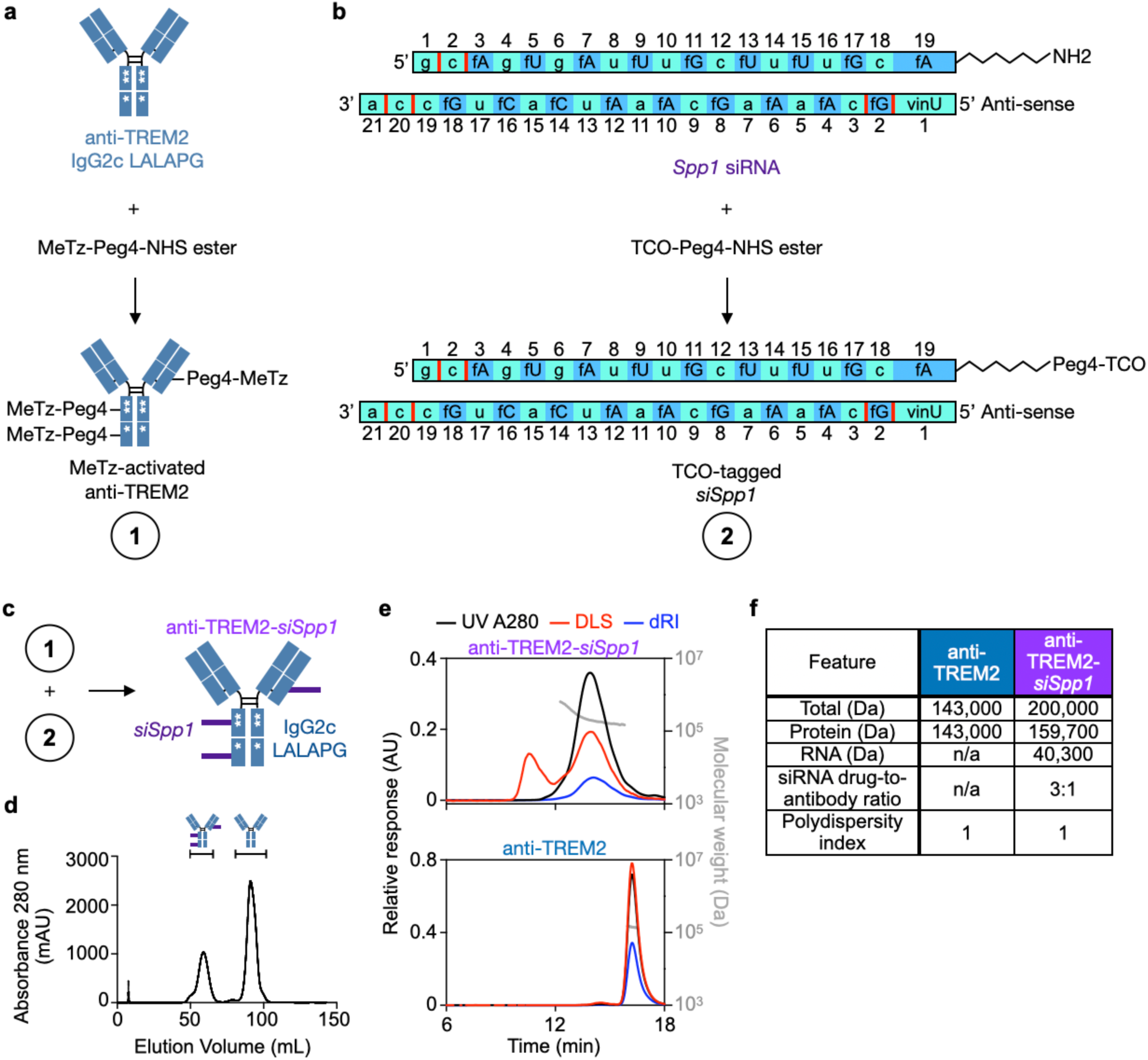
Antibody and siRNA conjugation. **a**, Method for functionalizing the anti-TREM2 antibody with MeTz-Peg4-NHS. **b**, Method for functionalizing the 3’ end of the siRNA sense strand with a TCO-Peg4-NHS moiety. fN = 2’-fluoro nucleotides; n = 2’-O-methyl nucleotides; vinU = 5’-E-VP uridine; red bars = phosphorothioate linkages. **c**, Click chemistry reaction scheme for conjugating the MeTz groups on the antibody with TCO-tagged siRNAs to form antibody-siRNA conjugates (ARCs). **d**, Size exclusion chromatography purification of ARC based on 280 nm absorbance. **e**, Representative size exclusion chromatography with multi-angle light scattering (SEC-MALS) of anti-TREM2 and anti-TREM2-*siSpp1* from two independent experiments. The protein concentration in the antibody sample or the conjugate was estimated from the differential refractive index increments (dn/dc), and the molecular mass was determined from the multi-angle light scattering (DLS) detector responses. **f**, Analysis of SEC-MALS to determine the siRNA drug-to-antibody conjugate ratio.

**Extended Data Fig. 5.**
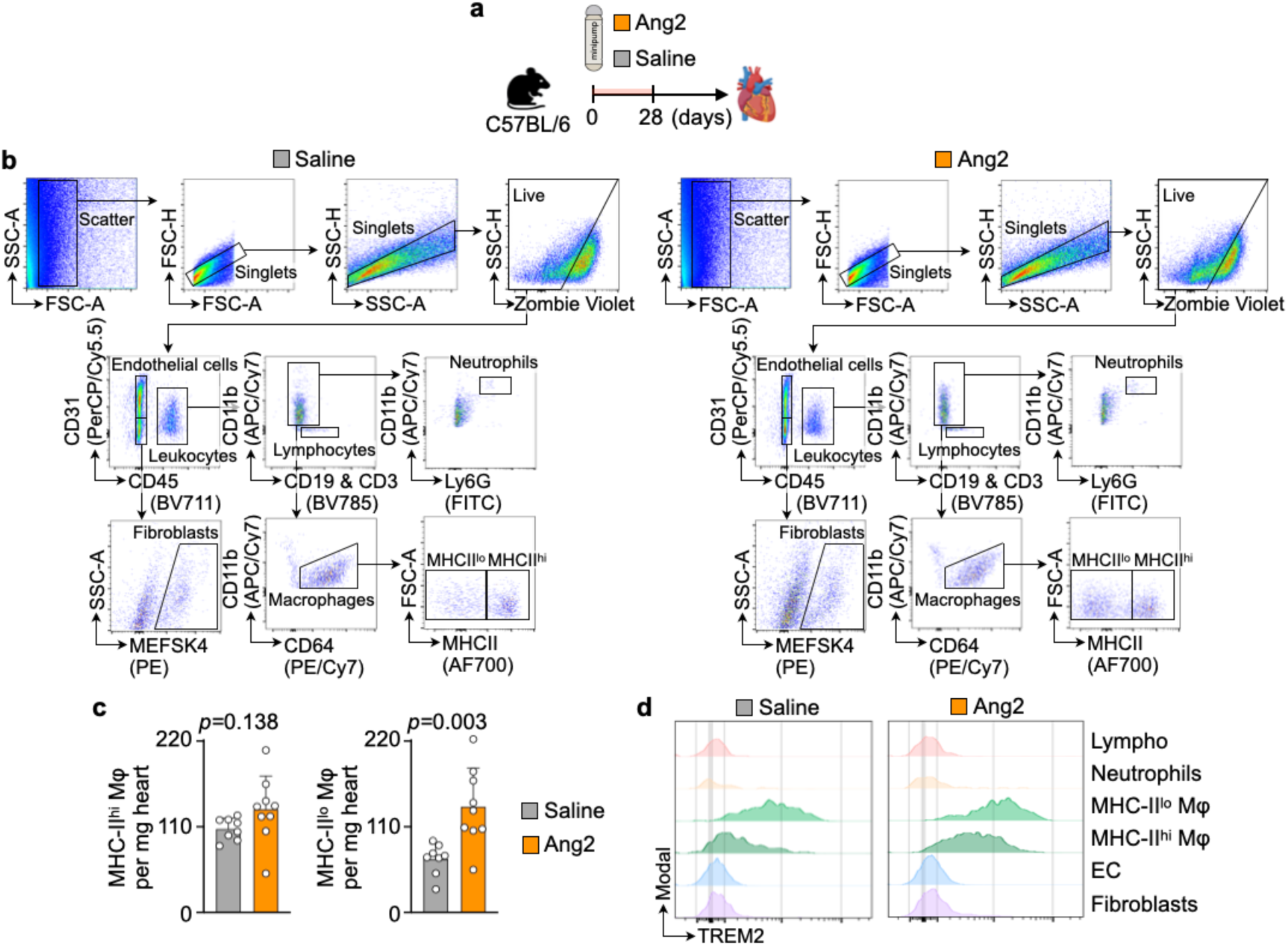
TREM2 expression on cardiac cells. **a**, Experimental outline of C57BL/6 mice infused with saline or Ang2 via minipumps for 28 days. **b**, Representative flow cytometry gating strategies to identify cell subsets in heart tissue from saline- and Ang2-treated mice. **c**, Enumeration of MHCII^hi^ and MHCII^lo^ macrophages in hearts from saline- and Ang2-treated mice by flow cytometry. Data are mean±s.d. with individual values for data distribution, n = 8 to 9 per group from two independent experiments, two-tailed Student’s t-test. **d**, Flow cytometry quantification of TREM2 on lymphocytes, neutrophils, MHCII^lo^ macrophages, MHCII^hi^ macrophages, endothelial cells, and fibroblasts in hearts from saline- and Ang2-treated mice.

**Extended Data Fig. 6.**
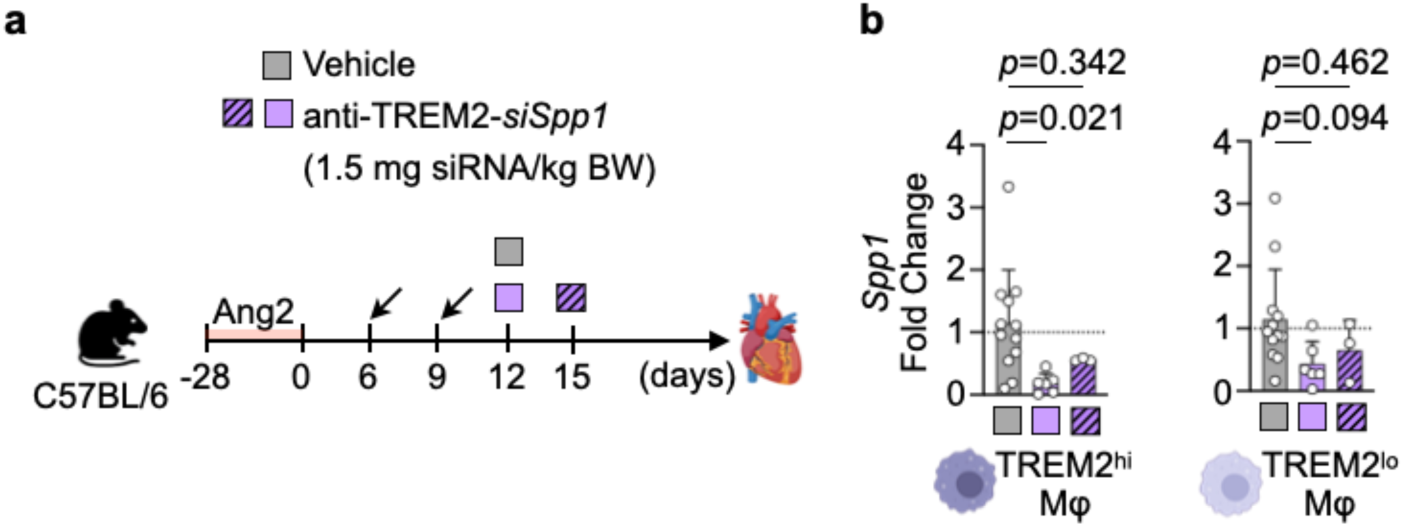
ARC silencing half-life is 6 days. **a**, Experimental outline of C57BL/6 mice infused with Ang2 via minipumps for 28 days followed by treatment with vehicle or anti-TREM2-*siSpp1* at 1.5 mg siRNA/kg body weight total dose equally divided between days 6 and 9. Hearts were excised either on day 12 or day 15. **b**, *Spp1* expression in TREM2^hi^ (left) and TREM2^lo^ (right) cardiac macrophages isolated on day 12 or 15 from anti-TREM2-*siSpp1*-treated hypertensive mice normalized to vehicle-treated hypertensive mice. Data are mean±s.d. with individual values for data distribution, n = 3 to 12 per group from four independent experiments, one-way ANOVA followed by Tukey’s multiple comparisons test.

**Extended Data Fig. 7.**
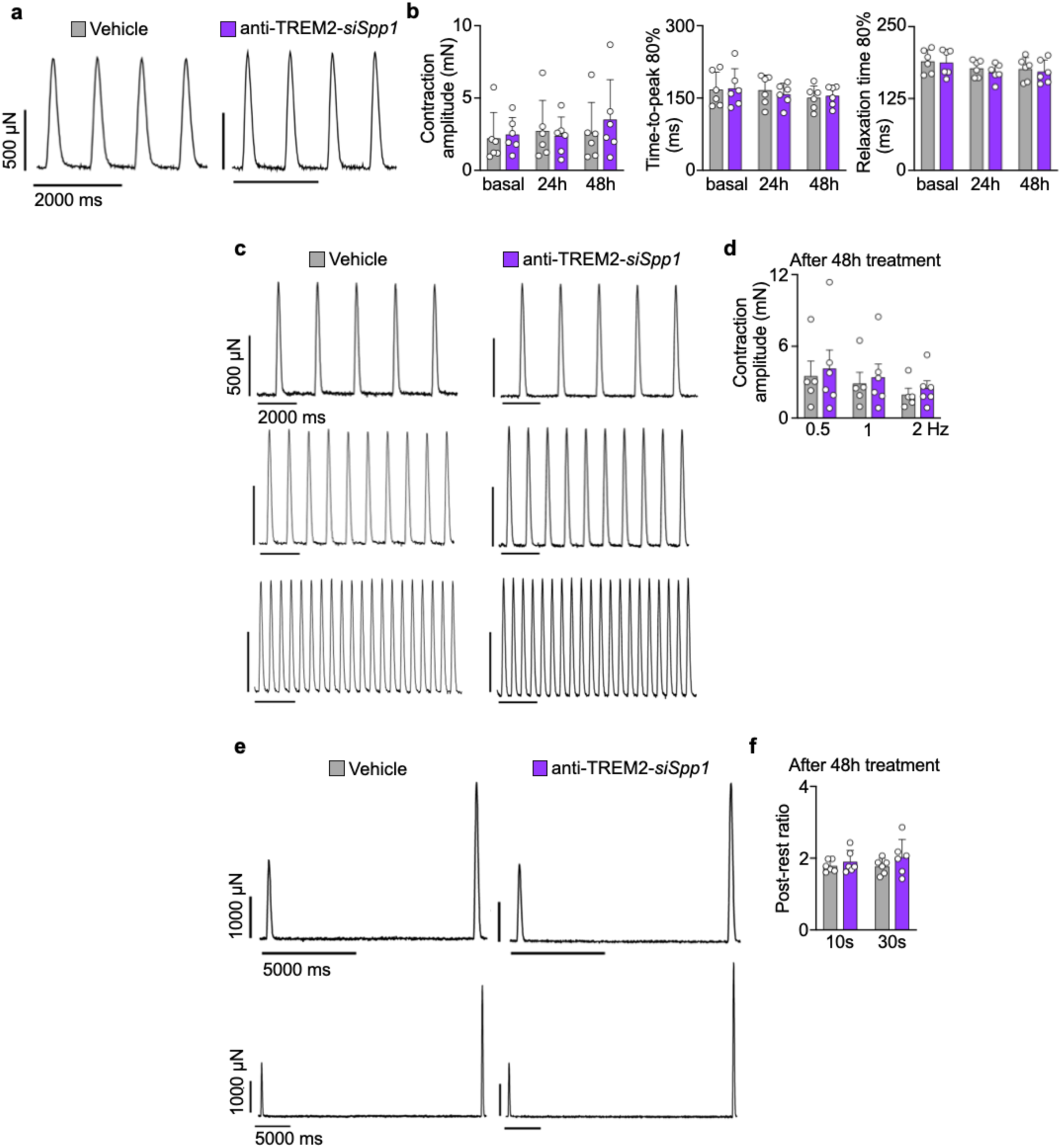
Contractility of ARC-treated human myocardium. **a**, Representative contraction traces of human myocardial slices 48 hours after treatment with either anti-TREM2-*siSpp1* or vehicle control. **b**, Quantification of contraction amplitude, time-to-peak 80% and relaxation time 80% of human myocardial slices treated with either anti-TREM2-*siSpp1* or vehicle control. n = 6 heart slices. **c**,**d**, Representative contraction traces (c) and contraction amplitudes (d) of vehicle- and anti-TREM2-*siSpp1*-treated myocardial slices during increasing stimulation rates (0.5, 1, and 2 Hz) testing the force-frequency relationship. n = 5 to 6 heart slices. **e**,**f**, Representative contractility traces (e) and post-rest ratios (f) of vehicle- and anti-TREM2-*siSpp1*-treated myocardial slices during stimulation pauses (10 s and 30 s) as a surrogate for sarcoplasmic reticulum function. n = 6 heart slices. Data are mean±s.d. with individual values for data distribution.

**Extended Data Fig. 8.**
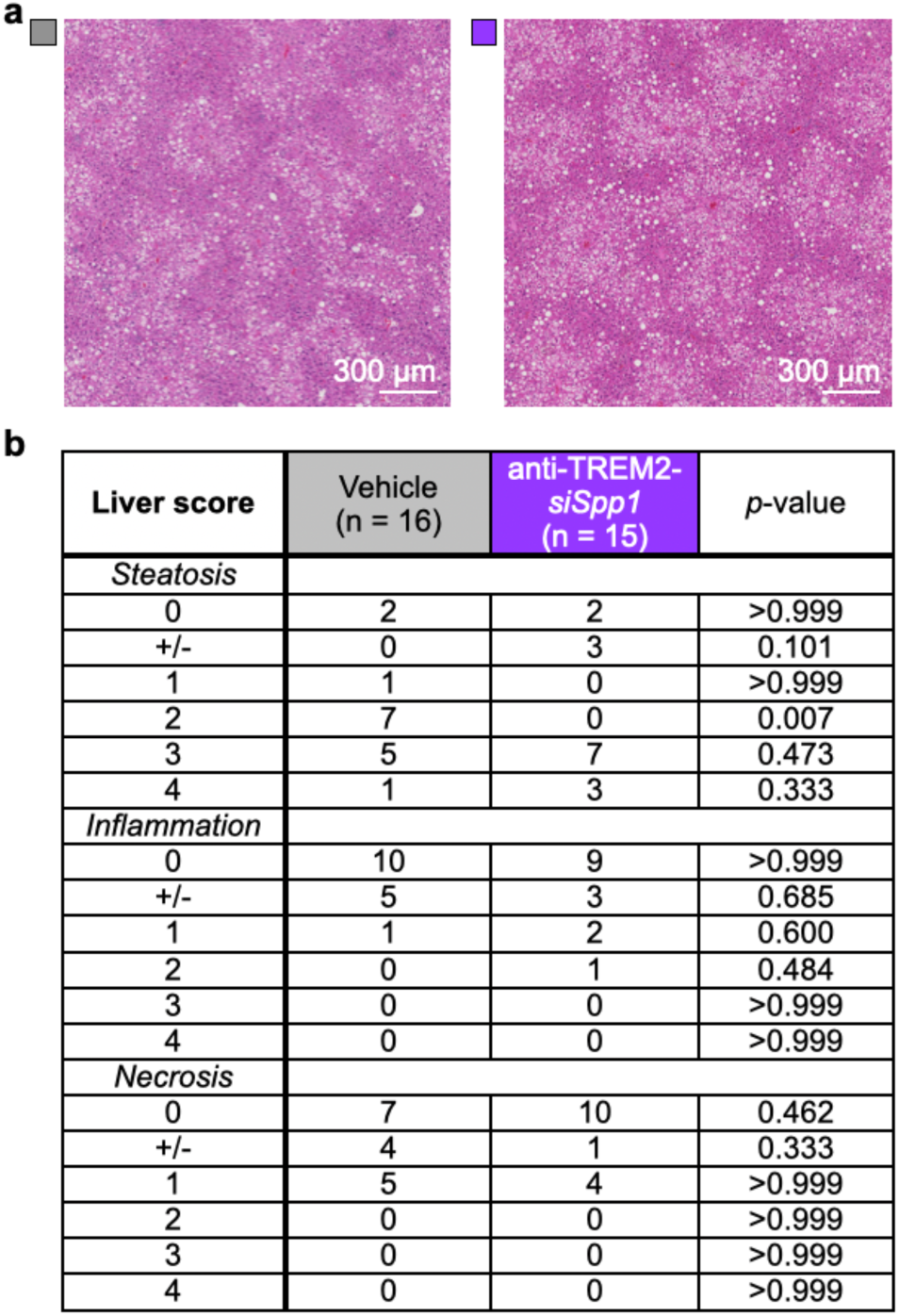
Liver histological features of treated HOMER mice. **a,** Representative H&E images of livers from vehicle- and anti-Trem2-*Spp1*-treated HOMER mice. Scale bars, 300 µm. **b**, Summary of histological features observed in liver sections from vehicle- and anti-Trem2-*Spp1*-treated HOMER mice. n = 15 to 16 per group from two independent experiments, Two-sided Fisher’s exact test. Scoring description: 0 = non; +/− = focal, minimal; 1 = mild; 2 = mild-moderate; 3 = moderate; 4 = severe.

**Extended Data Fig. 9.**
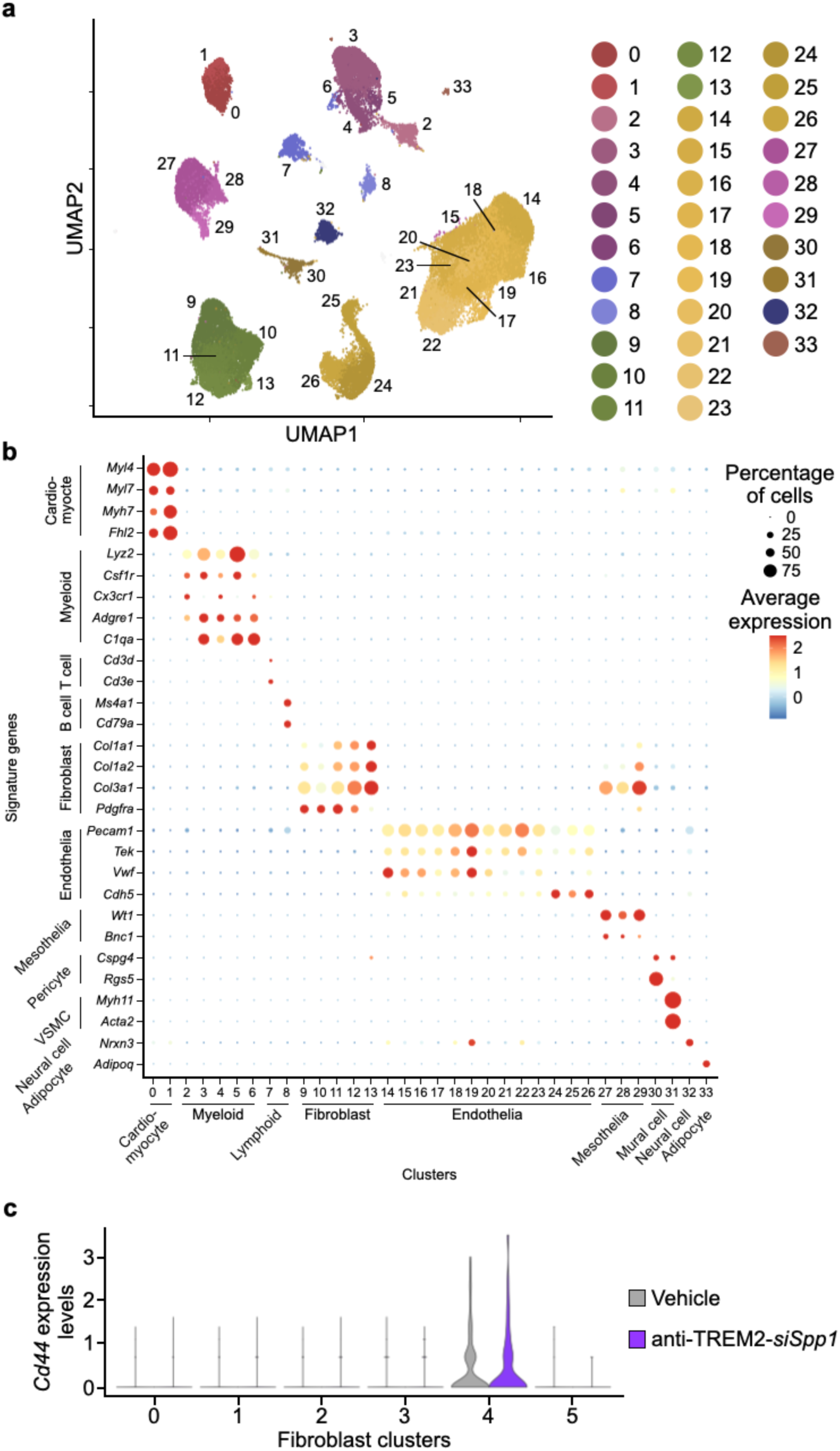
Annotating mouse left atrial nuclei. **a**, Clustering of 48,647 nuclei from left atrial tissues of three vehicle- and three anti-Trem2-*Spp1*-treated HOMER mice. **b**, Dot plot annotating nuclei clusters by signature gene expression. Color denotes z-score of average gene expression (red: high; blue: low); circle size indicates percentage of cells expressing the gene. **c**, *Cd44* expression levels in the 6 fibroblast clusters from three vehicle- and three anti-Trem2-*siSpp1*-treated HOMER mice.

